# Multi-lab EcoFAB study shows highly reproducible physiology and depletion of soil metabolites by a model grass

**DOI:** 10.1101/435818

**Authors:** Joelle Sasse, Josefine Kant, Benjamin J. Cole, Andrew P. Klein, Borjana Arsova, Pascal Schlaepfer, Jian Gao, Kyle Lewald, Kateryna Zhalnina, Suzanne Kosina, Benjamin P. Bowen, Daniel Treen, John Vogel, Axel Visel, Michelle Watt, Jeffery L. Dangl, Trent R. Northen

## Abstract

- There is a dynamic reciprocity between plants and their environment: On one hand, the physiochemical properties of soil influence plant morphology and metabolism, while on the other, root morphology and exudates shape the environment surrounding roots. Here, we investigate both of these aspects as well as the reproducibility of these responses across laboratories.
- The model grass *Brachypodium distachyon* was grown in phosphate-sufficient and phosphate-deficient mineral media, as well as in sterile soil extract, within fabricated ecosystem (EcoFAB) devices across four laboratories.
- Tissue weight and phosphate content, total root length, root tissue and exudate metabolic profiles were found to be consistent across laboratories and distinct between experimental treatments. Plants grown in soil extract were morphologically and metabolically distinct in all laboratories, with root hairs four times longer compared to other growth conditions. Further, plants depleted half of the investigated metabolites from the soil extract.
- To interact with their environment, plants not only adapt morphology and release complex metabolite mixtures; they also selectively deplete a range of soil-derived metabolites. The EcoFABs utilized here generated high inter-laboratory reproducibility, demonstrating that their value in standardized investigations of plant traits.

## Introduction

Plants adapt to their belowground environment by root morphological and metabolic plasticity. In turn, they influence soil physiochemical properties and root-associated organisms by creating the rhizosphere, an environmental niche formed by the physical structure of roots and the release of metabolites (root exudates). These complex root – environment interactions are challenging to study in general, and even more so in a manner that is reproducible across laboratories (Poorter *et al.*, 2012).

Root morphology and metabolism are affected by abiotic and biotic factors. Nutrient availability of soils for example can profoundly affect root morphology, and provoke changes in root metabolism. Phosphate limitation typically results in elongated lateral roots and root hairs in a context-dependent manner (Plaxton & Tran, 2011; Peret *et al.*, 2011; Nestler *et al.*, 2016), and increased exudation of organic acids that solubilize phosphate (Neumann & Martinoia, 2002; Plaxton & Tran, 2011; Thijs *et al.*, 2016). Root morphology and metabolism is further affected by microbes and microbial compounds (Venturi & Keel, 2016; Verbon & Liberman, 2016; Etalo *et al.*, 2018). The presence of plant growth-promoting bacteria can stimulate lateral root and root hair growth of Arabidopsis (López-Bucio *et al.*, 2007; Zamioudis *et al.*, 2013; Vacheron *et al.*, 2013). Plant responses to abiotic and biotic factors are likely intertwined, as illustrated recently by a study that linked phosphate stress in plants with the structure of root-associated microbial communities (Castrillo *et al.*, 2017). Thus, plant phenotypes in soil are a result of a complex response to abiotic and biotic factors, and an integrated view of root morphology and metabolism is necessary to gain a holistic understanding of plant – environment interactions.

Characterization of plant phenotypes in response to abiotic and biotic stresses in soil can have profound impact on agriculture, especially as many resources, such as phosphate-based fertilizers are limited (Cordell *et al.*, 2009), and global food demand is projected to have to increase by 60% by the year 2050 due to an ever-growing population (FAO. World food situation.)(). Grasses are central to biofuel production and provide 70% of human calories (Brutnell *et al.*, 2015). Thus, research on model grasses such as *Setaria viridis* and *Brachypodium distachyon* can inform growth strategies for many crops (Brutnell *et al.*, 2015). *B. distachyon* is gaining popularity as a model grass because of its small genome, short generation time, genetic tractability, and the availability of extensive germplasm and mutant collections (Hsia *et al.*, 2017). Additionally, since it uses C_3_ carbon fixation, it is a good laboratory model plant relevant to cereal crops such as barley, rice, and wheat. It has recently been utilized to investigate plant developmental processes, abiotic stresses, biotic interactions, and root morphology (Watt *et al.*, 2009; Brutnell *et al.*, 2015).

The relationship between plants and their environment is ideally studied in an agriculturally relevant field setting. Environmental factors, especially the type of soil in which plants are grown, are major determinants of root-associated microbial communities (Bulgarelli *et al.*, 2013; Edwards *et al.*, 2015), and of root morphology (Senga *et al.*, 2017). However, investigation of root morphology in soil is challenging due to its opacity, while investigation of exudation in soil is challenging due to its physiochemical complexity (Cai *et al.*, 2011). Specialized imaging techniques such as magnetic resonance imaging, computed tomography (Metzner *et al.*, 2015; Helliwell *et al.*, 2017), or the use of labeled plants (Rellán-Álvarez *et al.*, 2015) have been developed, but are not widely accessible or amenable to high throughput experimentation (Metzner *et al.*, 2015). Similarly, approaches for the investigation of root exudation in soils include the use of in-situ soil drainage systems (lysimeters) in fields (Strobel, 2001), which are low throughput and require complex installations, or of laboratory-based extraction methods that are based on flushing the soil with large volumes of liquids (Swenson *et al.*, 2015; Pétriacq *et al.*, 2017). Studying metabolites within rhizosphere soils is also challenging because of the complex mixture of plant- and microbe-derived metabolites, which are potentially altered by the chemistry and mineralogy of the soil investigated. A further challenge is the limited reproducibility of morphological and metabolic data generated (Massonnet *et al.*, 2010; Poorter *et al.*, 2012).

Due to these challenges in the field, root morphology and metabolism are often studied in laboratory settings. Laboratory environments can feature transparent substrates and mineral growth media devoid of complex chemical compounds present in soils, in order to allow straightforward investigation of plant traits. However, these highly artificial laboratory environments may not reproduce normal plant growth and plant-environment interactions that occur in the field. Thus, systems that allow the manipulation of aspects of natural systems in a controlled laboratory environment are desirable. Microfluidic devices are gradually improved to study for example heterogenous environments (Stanley *et al.*, 2017), and currently, these important devices are designed to accommodate plants with small roots such Arabidopsis for a growth period of several days to about two weeks (Parashar & Pandey, 2011; Jiang *et al.*, 2014; Stanley *et al.*, 2017). We recently reported on a modular growth system, the EcoFAB (Ecosystem Fabrication), which facilitates the evaluation of root morphology and exudation of various plants over the course of several plant developmental stages up to several weeks (Gao *et al.*, 2018). The EcoFAB design is purposely kept simple and inexpensive, to allow for straightforward design and manufacturing of EcoFABs for various experimental questions. The EcoFABs also address the challenge of studying plant growth in various environments, such as chemically simple or complex hydroponic setups, including the ability to add solid substrates such as sand or soil. In addition, microbes can be added to EcoFAB chambers, and the system is compatible with chemical imaging (Gao *et al.*, 2018). One of the key distinctions of a standardized system such as the EcoFAB is the reproducibility of data generated.

The study presented here aimed to test the reproducibility of EcoFABs across multiple laboratories in assessing the response of the model grass *B. distachyon* in different growth media. Phosphate-sufficient and - deficient mineral media were chosen to assess the performance of the EcoFAB system in reproducing well-described effects of phosphate starvation, and a complex sterilized soil extract was chosen as representation of a more natural environment with yet uncharacterized effects on plant morphology and metabolism. We hypothesized that the use of the EcoFAB system produces data reproducible across laboratories, and that *B. distachyon* grown in the various media would result in distinct metabolic and morphological changes.

## Material and Methods

### EcoFAB preparation

EcoFAB devices were fabricated according to the published method (Gao *et al.*, 2018). Briefly, an 1:10 silicone elastomer curing agent : base mixture (PDMS, Ellsworth Adhesives) was poured onto a 3D-printed mold, and allowed to solidify at 80°C for 4 h. The PDMS layer was separated from the mold, the edged trimmed, and permanently bonded to a glass microscope slide. The EcoFAB device and outer chamber were sterilized by incubation in 70% v/v ethanol for 30 min, followed by incubation in 100% v/v ethanol for 5 min. After evaporation of residual ethanol, the EcoFAB device was rinsed three times with the growth medium of choice before transferring seedlings.

### Plant growth conditions

All experiments were performed with Brachypodium distachyon Bd21-3 (Vogel & Hill, 2007). Seeds were dehusked and sterilized in 70% v/v ethanol for 30 s, and in 6% v/v NaOCl, 0.1% v/v Triton X-100 for 5 min, followed by five wash steps in water. Seedlings were germinated on 0.5x Murashige & Skoog plates (2.2 g l^−1^ MS medium, MSP01, Caisson Laboratories with 1650 mg L^−1^ NH_4_NO_3_, 6.2 mg L^−1^ H_3_BO_3_, 332.2 ml L^−1^ CaCl_2_, 0.025 ml L^−1^ CoCl_2_, 0.025 ml L^−1^ CuSO_4_, 37.26 mg L^−1^ C_10_H_14_N_2_Na_2_O_8_, 27.8 mg L^−1^ FeSO_4_*7H_2_O, 180.7 mg L^−1^ MgSO_4_, 16.9 ml L^−1^ MnSO_4_*H_2_O, 0.25 mg L^−1^ NaMoO_4_*2H_2_O, 0.83 mg L^−1^ KI, 1900 mg L^−1^ KNO_3_, 170 mg L^−1^ KH_2_PO_4_, 8.6 mg L^−1^ ZnSO_4_*7H_2_O; 6% w/v Bioworld Phytoagar, 401000721, Fisher Scientific, pH adjusted to 5.7) in a 16 h light / 8 h dark regime at 24°C. EcoFABs were sterilized as published, and seedlings transferred to EcoFAB chambers at three days after germination (dag) as previously described (Gao *et al.*, 2018). Seedlings with comparable size were picked to conduct the experiment, and were distributed in a random manner to the various EcoFABs. EcoFABs were incubated in a 16 h light / 8 h dark regime at 24°C, with 150 μE illumination. The EcoFABs were filled with 2 ml of 0.5x MS (*B. distachyon* grows without phenotypically detectable nutrient limitation, ‘phosphate-sufficient’, 2.2 g l^−1^ MS medium, MSP01, Caisson Laboratories, pH adjusted to 5.7), 0.5x MS-P (B distachyon leaves turn yellow as a sign of malnutrition, ‘phosphate-deficient’, 2.2 g l^−1^ MS medium without phosphate, composition is the same as MSP01 without 170 mg L^−1^ KH_2_PO_4_, MSP11, Caisson Laboratories, pH adjusted to 5.7), or soil extract. The soil extract was prepared by incubating 100 g of a standard greenhouse soil (Pro-Mix PGX, Hummert International) in 1 l of water for 16 h at 4°C and gentle shaking, followed by filtration through a 0.2 μm cellulose nitrate filter (09-761-104, Corning) for sterilization. The soil extract was stored at 4°C, and its phosphate content was determined as 145 μM (Ames, 1966), which is four times lower compared to 0.5x MS. Although we did not perform additional nutrient analyses, it is likely that levels of other nutrients besides phosphate are also low, compared to 0.5x MS.

A comparative study of *B. distachyon* in EcoFABs versus plates was performed by laboratory 1, in which *B. distachyon* seeds were sterilized and germinated on 0.5x MS plates for 3 days as described above, then either transferred to EcoFAB growth chambers containing 0.5x MS liquid medium as described (Gao *et al.*, 2018), or to 0.5x MS phytoagar plates. Roots were imaged weekly, and total root area was measured using the Image J software suite (version 2.0.0). For the developmental timecourse, plants were grown in EcoFAB chambers in 0.5x MS for up to 43 days, and exudates were collected at indicated times (Fig. S1), frozen, and stored at −80°C. Metabolites were analyzed as described below.

### EcoFAB inter-laboratory experiment

An overview of the experimental procedure is provided in Fig. 1, and the participating laboratories are listed in Table S1. The following material was distributed from laboratory 1 to the participating laboratories: EcoFAB growth chambers, micropore tape to seal the EcoFABs, *B. distachyon* seeds, MS powder, MS-P powder, liquid soil extract (see ‘Plant growth conditions’), phytoagar, light & temperature data loggers (HOBO Onset, UA-002-08), and a detailed protocol for plant growth and experimental procedures. The experiments were conducted in parallel by the different laboratories. Each participating laboratory sterilized EcoFABs and seeds as described (Gao *et al.*, 2018). Growth conditions were monitored throughout the experiment, and are reported in Table S1. Plants were grown in quadruplicates for each experimental condition, and one control EcoFAB was set up per condition without plants. Sterility was monitored throughout the experiment by plating 50 μl of growth media on Luria-Bertani (LB) plates every week. Contaminated chambers were excluded from analysis.

**Fig. 1.**
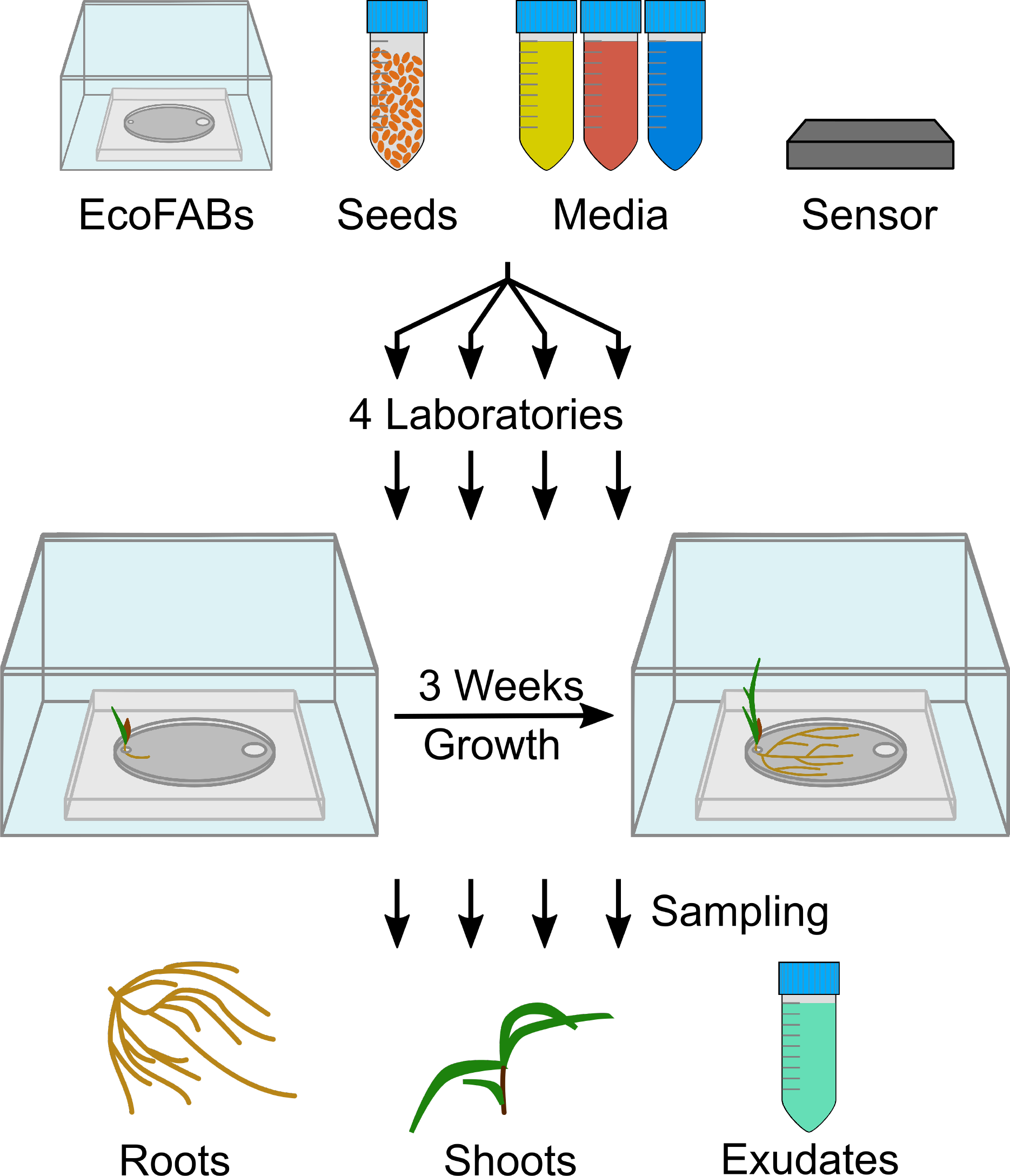
Experimental setup of the reproducibility experiment. Illustration of the reproducibility experiment: EcoFABs, *B. distachyon* seeds, growth media (0.5x MS: yellow, 0.5x MS-P: red, soil extract: blue), and light/temperature sensors were distributed to the participating laboratories. Each laboratory germinated the seeds, transferred seedlings to sterilized EcoFABs, and grew the plants for 21 days. Root and shoot tissue as well as root exudates were sampled for downstream analysis.

Root systems in EcoFABs were imaged at 7, 14, and 21 days after transfer (dat) to the EcoFAB chambers. Total root length was quantified by laboratory 1 with the SmartRoot plugin (version 4.21) for the ImageJ software (version 2.0.0)(Lobet *et al.*, 2011). Root hairs were imaged at 21 days with 10x magnification, and their length was determined with ImageJ. The data presented is an average of three measurements per imaged root.

Growth media was replenished to 2 ml three times a week, and the media were exchanged fully at 20 dat. This medium was removed through the sampling port by pipetting after 24 h of further incubation, and the volume was recorded. The root exudates were frozen immediately, stored at −80°C, and shipped to laboratory 1 for metabolite analysis. The fresh weight of root and shoot tissue was recorded, and the tissue was immediately frozen and stored at −80°C. The tissue was homogenized by the participating laboratories by their method of choice (mortar and pestle with liquid nitrogen, or steel beads with a bead beater). An aliquot of the tissue was utilized for phosphate content determination by all participating laboratories (Ames, 1966), and an aliquot was sent back to laboratory 1 for metabolite analysis.

### Liquid chromatography mass spectrometry sample extraction

Homogenized root tissues were extracted two times with 700 μl 100% LC/MS grade methanol (CAS 67-56-1, Honeywell Burdick & Jackson, Morristown, NJ) for 1 h at 4°C. The samples were centrifuged for 5 min at 5000 g, 4°C, supernatants were pooled and evaporated under vacuum at 25°C until dry. The samples were resuspended in 100% LC/MS grade methanol with 15 μM internal standards (767964, Sigma-Aldrich) with a volume relative to the sample fresh weight (11 mg / 100 μl).

Frozen root exudates were lyophilized using a Labconco FreeZone lyophilizer, resuspended in 500 μl LC/MS grade methanol (CAS 67-56-1, Honeywell Burdick & Jackson, Morristown, NJ), sonicated for 15 min in a water bath at 23°C, and incubated at 4°C for 16 h for salt precipitation. Samples were then centrifuged for 5 min at 5000 g, 4°C, supernatants were transferred to new microcentrifuge tubes, and evaporated at 25°C under vacuum until dry. Samples were resuspended in 100% LC/MS grade methanol with 15 μM internal standards (767964, Sigma-Aldrich) with a volume relative to the root tissue fresh weight, and the root exudate volume collected (20 μl methanol 100 mg-1 fresh weight ml-1 exudate volume).

### Liquid chromatography – mass spectrometry method and analysis

Metabolites in samples were chromatographically separated using hydrophilic liquid interaction chromatography on a SeQuant 5 μm, 150 × 2.1 mm, 200 Å zic-HILIC column (1.50454.0001, Millipore) and detected with a Q Exactive Hybrid Quadrupole-Orbitrap Mass Spectrometer equipped with a HESI-II source probe (ThermoFisher Scientific). For chromatographic separations, an Agilent 1290 series HPLC system was used with a column temperature of 40°C, 3 μl sample injections, and 4°C sample storage. A gradient of mobile phase A (5 mM ammonium acetate in water) and B (5 mM ammonium acetate, 95% v/v acetonitrile in water) was used for metabolite retention and elution as follows: column equilibration at 0.45 ml min^−1^ in 100% B for 1.5 min, a linear gradient at 0.45 ml min^−1^ to 35% A over 13.5 minutes, a linear gradient to 0.6 ml min^−1^ and to 100% A over 3 min, a hold at 0.6 ml min^−1^ and 100% A for 5 min followed by a linear gradient to 0.45 ml min-1 and 100% B over 2 min and re-equilibration for an additional 7 min. Each sample was injected twice: once for analysis in positive ion mode and once for analysis in negative ion mode. The mass spectrometer source was set with a sheath gas flow of 55, aux gas flow of 20 and sweep gas flow of 2 (arbitrary units), spray voltage of |±3| kV, and capillary temperature of 400°C. Ion detection was performed using the Q Exactive’s data dependent MS2 Top2 method, with the two highest abundance precursory ions (2.0 m/z isolation window, 17,500 resolution, 1e5 AGC target, 2.0m/z isolation window, stepped normalized collisions energies of 10, 20 and 30 eV) selected from a full MS pre-scan (70-1050 m/z, 70,000 resolution, 3e6 AGC target, 100 ms maximum ion transmission) with dd settings at 1e3 minimum AGC target, charges excluded above |3| and a 10 s dynamic exclusion window. Internal and external standards were included for quality control purposes, with blank injections between every unique sample.

### Metabolite identification and statistical analysis

LC/MS data was analyzed with Metabolite Atlas to construct extracted ion chromatograms corresponding to metabolites contained within our in-house standards library (https://github.com/biorack/metatlas)(Bowen & Northen, 2010; Yao *et al.*, 2015). For metabolite identification, chemical classes were assigned using the ClassyFire compound classification system (Djoumbou Feunang *et al.*, 2016). Metabolites were identified following the conventions defined by the Metabolomics Standards Initiative (Sumner *et al.*, 2007)(Table S2, S3). All assignments were of the highest confidence (‘level 1’ MSI identifications), which is identified as at least two orthogonal measures vs. authentic chemical standards (e.g. retention time and fragmentation spectra). In all cases we used three orthogonal measures, retention time (within 1 minutes vs. standard), fragmentation spectra (manual inspection), and accurate mass (within 20 ppm). In general accurate masses were within 5 ppm, though the error was higher for low mass ions in negative mode. Peak height and retention time consistency for the LC/MS run was ascertained by analyzing quality control samples that were included at the beginning, during, and at the end of the run. Internal standards were used to assess sample-to-sample consistency for peak area and retention times.

Metabolite background signals detected in the extraction blanks, 0.5x MS and 0.5x MS-P control samples were subtracted from the experimental sample peak heights. Further, metabolite peak heights were normalized by setting the maximum peak height detected in any sample to 100%. The method utilized here allows for the relative comparison of peak heights between samples (e.g. if a compound of interest is present in significantly different amounts between samples), but not for absolute metabolite level quantification (e.g. μg of a compounds of interest per gram tissue). To explore the variation between growth conditions, the metabolite profiles were PCA-ordinated, and the 95% confidence level was displayed as ellipses for each treatment. Hierarchical clustering analysis with a Bray Curtis Dissimilarity Matrix was performed with the python 2.7 Seaborn package. The significance between root tissue as well as root exudate metabolic profiles was analyzed with the python SciPy ANOVA test coupled to a python Tukey’s honestly significant difference test with alpha = 0.05 corresponding to a 95% confidence level for each metabolite. Statistically significant metabolites were displayed as bar graphs, where the sum of all values added up to 100% (Fig. 4, Fig. S4), or as fold change for soil extract exudates divided by soil extract controls (Fig. 5).

## Results

### The EcoFAB growth system design and benchmarking

The EcoFAB device is comprised of a PDMS layer bonded to a glass slide, and an outer box to maintain sterility (Gao *et al.*, 2018) with a plant reservoir to hold the seedling, and a sampling port for to addition or exchange of growth medium (Fig. S1A). *B. distachyon* can be grown in EcoFABs for multiple weeks (Fig. S1B depicts a three-week-old *B. distachyon* plant), facilitating the investigation of various developmental stages from seedlings to adult plants.

We benchmarked *B. distachyon* growth in the EcoFAB vs. on standard agar plates. We found that *B. distachyon* roots develop similarly in EcoFABs containing 0.5x MS medium as compared to growth on 0.5x MS agar plates over the course of five weeks, with no significant differences in total root area observed except for week 2 (p = 0.05) (Fig. S1C). In addition, sampling of *B. distachyon* root exudates at different developmental stages showed a gradual shift of exudate profiles over time (Fig. S1D), consistent with reports for plants in other growth systems (Chaparro *et al.*, 2013; Zhalnina *et al.*, 2018).

### Multi-lab investigation of EcoFAB data reproducibility

EcoFab materials were distributed to four participating laboratories that ran the same experiment in parallel, investigating morphological and metabolic changes of *B. distachyon* grown in phosphate-sufficient, phosphate-deficient, or soil extract medium (4.3 times less phosphate than phosphate-sufficient medium). Roots were imaged on a weekly basis, and after three weeks, each laboratory determined the fresh weight and phosphate content of root and shoot tissue, and sampled root tissue and exudates for LC/MS analysis (Fig. 1).

Growth conditions (light intensity, day length, and temperature) were comparable between laboratories throughout the experiment (Table S1). The fresh weight and phosphate content were consistent across laboratories, and different between experimental treatments (Fig. 2a, b): as expected, phosphate-deficient plants had significantly lower phosphate content, and less than half the fresh weight of phosphate-sufficient plants (Tukey’s test, p=0.05). Interestingly, soil extract-grown plants showed a mixed response, in that they resembled phosphate-deficient plants in phosphate content and shoot weight, but their root weight was significantly lower than of phosphate-sufficient plants, and more similar to phosphate-sufficient plants. The root:shoot fresh weight ratio averaged across all laboratories was 0.9 for phosphate-sufficient plants, 1.3 for phosphate-deficient plants, and 1.8 for soil extract-grown plants (Fig. S2).

**Fig. 2.**
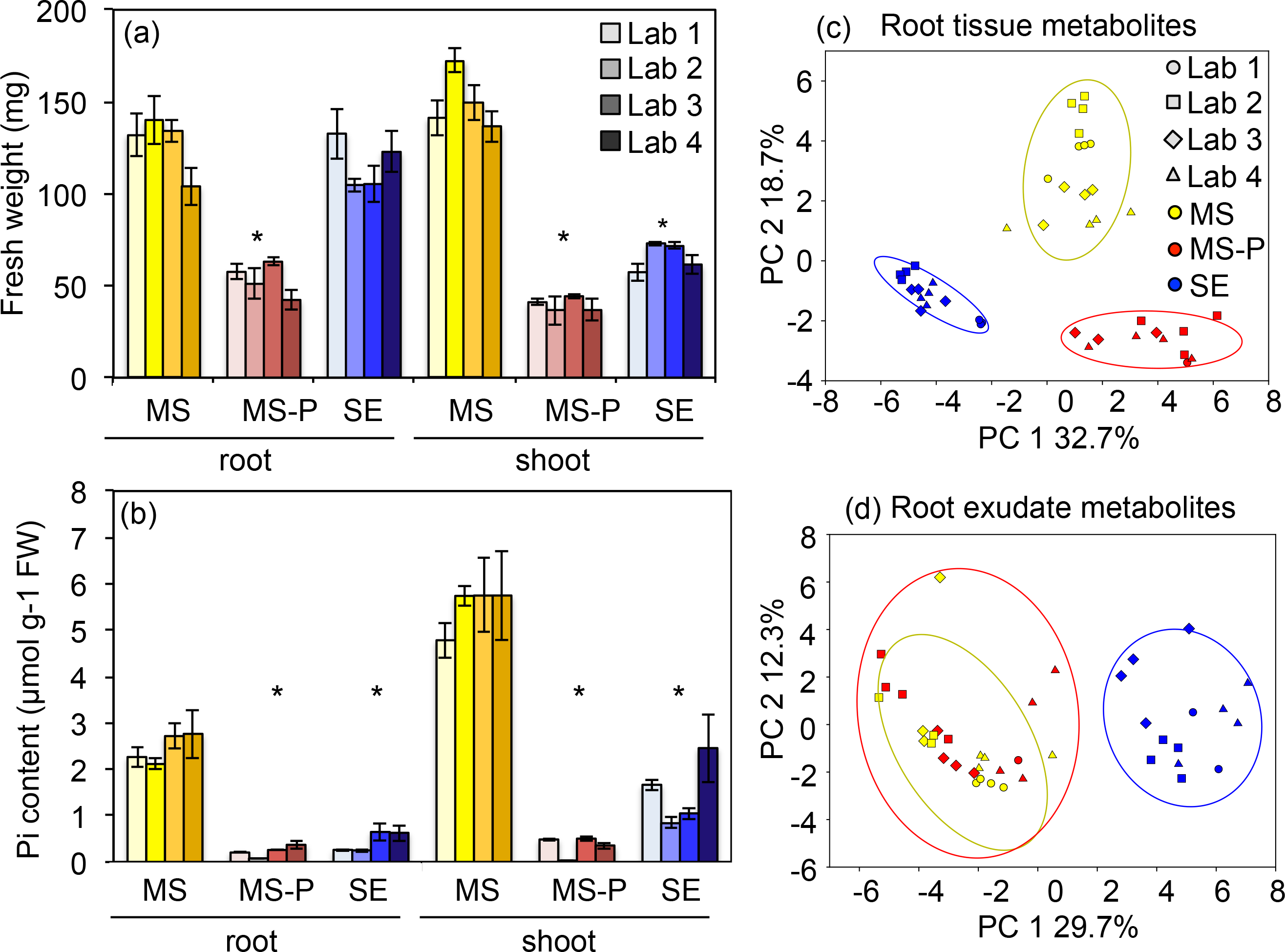
Inter-laboratory morphological and metabolic consistency. *B. distachyon* was grown in 0.5x MS (MS, yellow), 0.5x MS-P (MS-P, red), or soil extract (SE, blue) for three weeks. Root and shoot fresh weight (a) and phosphate content (b) were determined by the participating laboratories. Data are means ± s.e.m. (N > 9). Asterisks indicate significant differences between experimental treatments (Anova, p < 0.05). Principal component analysis of normalized peak heights of ground root tissue metabolites (c) and root exudate metabolites (d). Hierarchical clustering for the metabolite data is shown in Fig. S3. PC, principal component.

Upon receiving samples from each laboratory following the experiment, laboratory 1 extracted and analyzed metabolites, generating metabolite profiles from root tissues and exudates using LC/MS. The metabolic profiles of root tissues were comparable across laboratories, and reproducibly demonstrated a clear separation between experimental conditions in a principal component analysis plot and in a hierarchical clustering analysis (Fig. 2c, Fig. S3a, Tukey’ honestly significance test, p = 0.05). Similarly, the metabolic profiles of root exudates were comparable across laboratories and showed a separation between soil extract and other growth conditions (Fig. 2d, Fig. S3b).

Root morphology (quantified by laboratory 1) was similarly different between experimental treatments. Plants grown in phosphate-sufficient conditions formed root systems extending across most of the EcoFAB root chambers, whereas phosphate-deficient roots did not reach as far. Soil-extract grown roots also reached across the entire root chamber, with overall less roots compared to phosphate-sufficient plants, but visibly elongated root hairs (Fig. 3a). Quantification of total root length averaged across laboratories was 7 cm at 7 days after transfer (dat) for all plants, increased to 40 cm, 22 cm, and 30 cm at 14 dat, and further to 114 cm, 48 cm, and 67 cm for phosphate-sufficient, phosphate-deficient, and soil extract-grown plants, respectively. Differences between experimental treatments were first visible 14 dat with phosphate-deficient plants exhibiting shorter total root length than phosphate-sufficient plants (Tukey, p=0.05), but became more pronounced by 21 dat, with phosphate-sufficient plants exhibiting longer total root length than those grown in soil extract, which in turn were longer than of phosphate-deficient plants (Fig. 3b). Interestingly, root morphology varied somewhat between laboratories, the absolute measurements differed up to a factor of 2, with plants grown in laboratories 1 and 4 exhibiting consistently higher total root length than plants of laboratories 2 and 3 (Fig. S4). Specifically, total root length was 75-150 cm in phosphate-sufficient, 32-62 cm in phosphate-deficient, and 44-87 cm in soil extract conditions (Fig. S4).

**Fig. 3.**
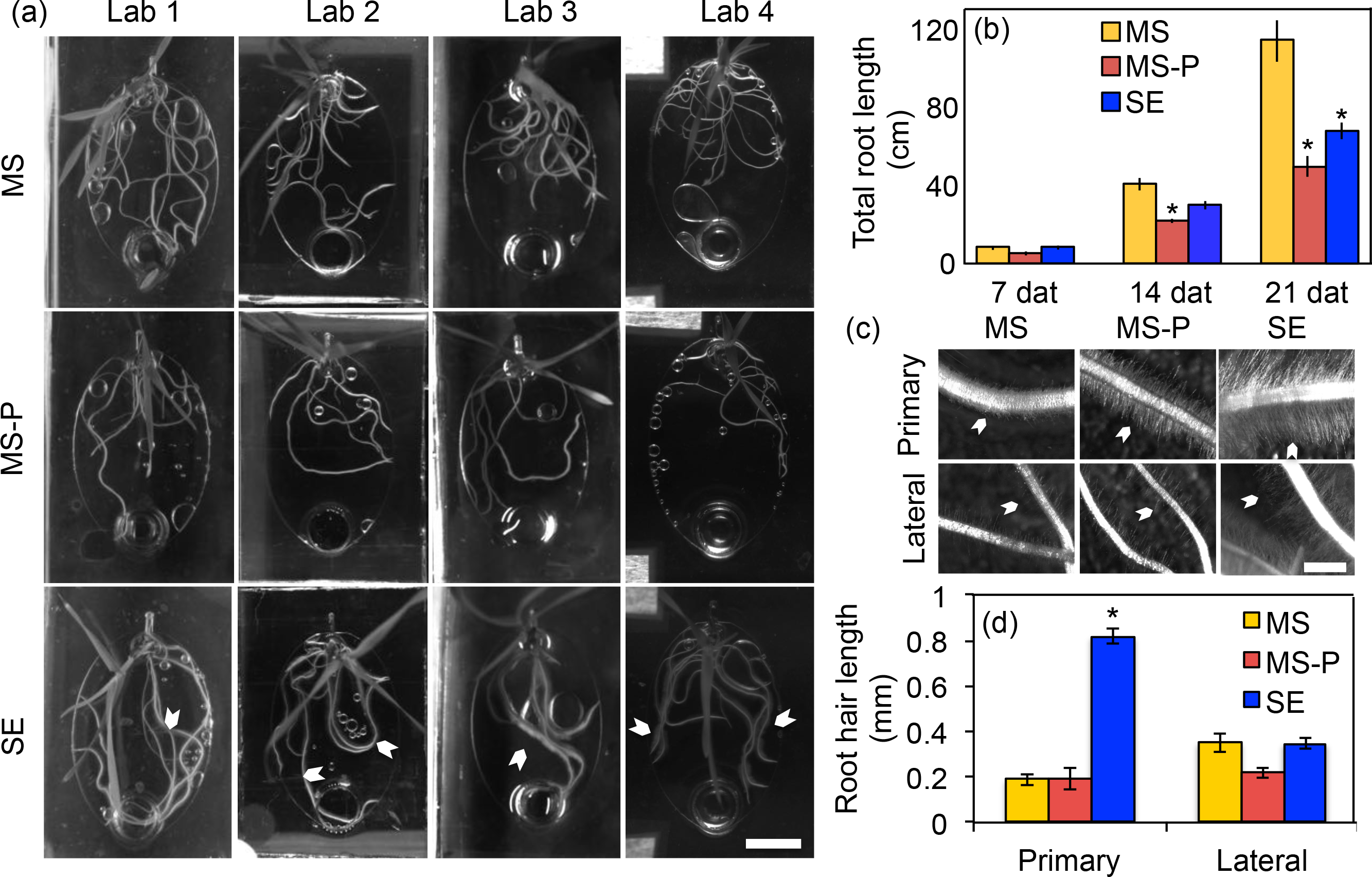
Root morphology. (a) Representative pictures of 14 dat (days after transfer) *B. distachyon* in EcoFAB chambers in 0.5x MS (MS), 0.5x MS-P (MS-P), or soil extract (SE) for the different laboratories (Lab1-4). Note the long root hairs in soil-extract growing plants (arrowheads). Brightness and contrast were adjusted for better display. Scale bar: 1 cm. (b) Total root length 7, 14, and 21 dat averaged across laboratories. The same data is displayed per lab in Fig. S4. Data are means ± s.e.m. (N > 9). (c) Root hair morphology. Arrowheads point to root hairs. Scale bar = 1 mm. (d) Root hair length at 21 dat for primary and lateral roots. Data are means ± s.e.m. (N > 9). Asterisks indicate significant differences within a group of bars (Anova, p < 0.05).

To summarize, the root and shoot fresh weight and phosphate content, root and exudate metabolic profiles, and total root length was consistent across laboratories and distinct for the experimental treatments.

### Distinct root morphology in soil extract

In addition to the high root:shoot ratio observed for soil extract-grown plants (Fig. S2), plants grown in soil extract had longer root hairs compared to plants grown in other conditions which were visible even under low-magnification (Fig. 3a, c). Interestingly, quantification revealed that root hairs on primary soil extract-grown roots reached a length of 0.8 mm, which was four times longer compared to phosphate-sufficient or phosphate-deficient grown roots. Root hair length of lateral roots remained unchanged (Fig. 3d).

### Metabolic analysis of root tissue and exudates

Metabolites extracted from root tissue and root exudates were found to be distinct between experimental treatments (Fig. 2c, d). Based on authentic metabolite standards, a broad range of metabolites was detected in root tissues as well as in exudates, among them organic acids, carbohydrates, nucleosides/nucleotides/nucleic bases, amino acids and other nitrogenous compounds, benzenoids, and fatty acids.

Half of the metabolites detected in root tissue extracts (52 out of 117 compounds) were significantly different in pairwise comparisons of experimental treatments, with 28% having highest abundance in phosphate-sufficient, 30% in phosphate-deficient, and 25% in soil extract-grown roots (Fig. 4, Table S2). The significantly different metabolites (p-value < 0.05) could be grouped into four main clusters (Fig. 4): Cluster I consists of three metabolites significantly different between all experimental treatments. Cluster II is composed of metabolites abundant in phosphate-sufficient roots, including: nucleosides, organic acids, amino acids, and notably, all phosphorous compounds present in this dataset. The higher abundance of phosphorous compounds in phosphate-sufficient roots compared to phosphate-deficient or soil extract-grown roots is in line with the phosphate quantification of plant tissues (Fig. 2b), in which highest free phosphate was detected in phosphate-sufficient plants, as would be expected. Cluster III includes metabolites abundant in phosphate-deficient roots. All these metabolites are nitrogenous compounds, likely due to the nitrogen-phosphate imbalance of phosphate-deficient plants. Cluster IV contains metabolites distinct for soil extract-grown roots, and is split in two subclusters. IVa includes metabolites with low abundance in soil extract roots, which are mostly nitrogenous compounds, whereas IVb includes metabolites with high abundance in soil extract roots, which are mostly organic acids.

**Fig. 4.**
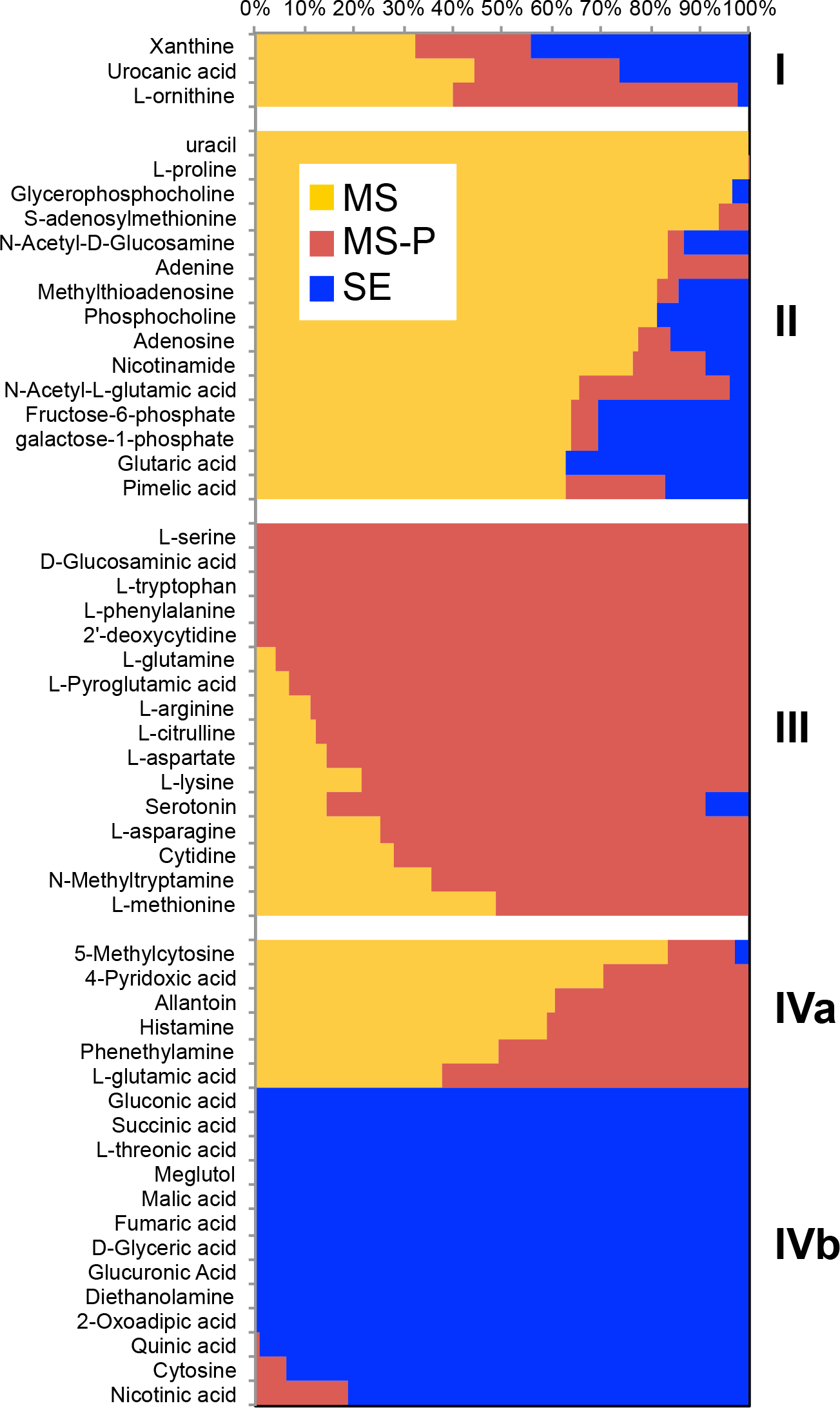
Characteristic metabolites detected in different root tissues. Normalized relative peak height of metabolites differing between roots grown in 0.5x MS (MS, yellow), 0.5x MS-P(MS-P, red), and soil extract (SE, blue) (Anova, p < 0.05). Metabolite cluster are indicated by roman numerals.

Overall, 137 metabolites were identified in root exudates (Table S3). Only phenylacetaldehyde was significantly different between exudates of phosphate-sufficient and – deficient plants (Table S3), which explains why these conditions are not separated in a principal component analysis (Fig. 2d). Plants grown in soil extracts had a distinct exudate composition with 27 and 25 distinct compounds vs. phosphate-sufficient and phosphate-deficient root exudates, respectively. Most of these distinct metabolites were most abundant in soil extract controls (no plant), showed medium abundance in soil extract exudates, and had low abundance in the other conditions (Fig S4).

Metabolite comparisons between soil extract with and without plants revealed that half of the metabolites detected (74 of 136 compounds) were altered in abundance, causing a distinct grouping in a principal component analysis (Fig. S6). Fifty percent of these metabolites were depleted in the presence of plants (Table S3, Fig. S5). Although individual metabolite levels varied somewhat across laboratories, this finding was consistent across participating laboratories (Fig. S7). Distinct metabolites included organic acids, carbohydrates, amino acids, and nucleosides, and these compounds contain various groups such as phosphate, nitrogen, or sulfur (Fig. 5, Table S3). Furthermore, citric acid exhibited an interesting but statistically insignificant trend of higher abundance in soil extract exudates vs. controls (Table S3, Anova p = 0.23, T-Test p = 0.04).

**Fig. 5.**
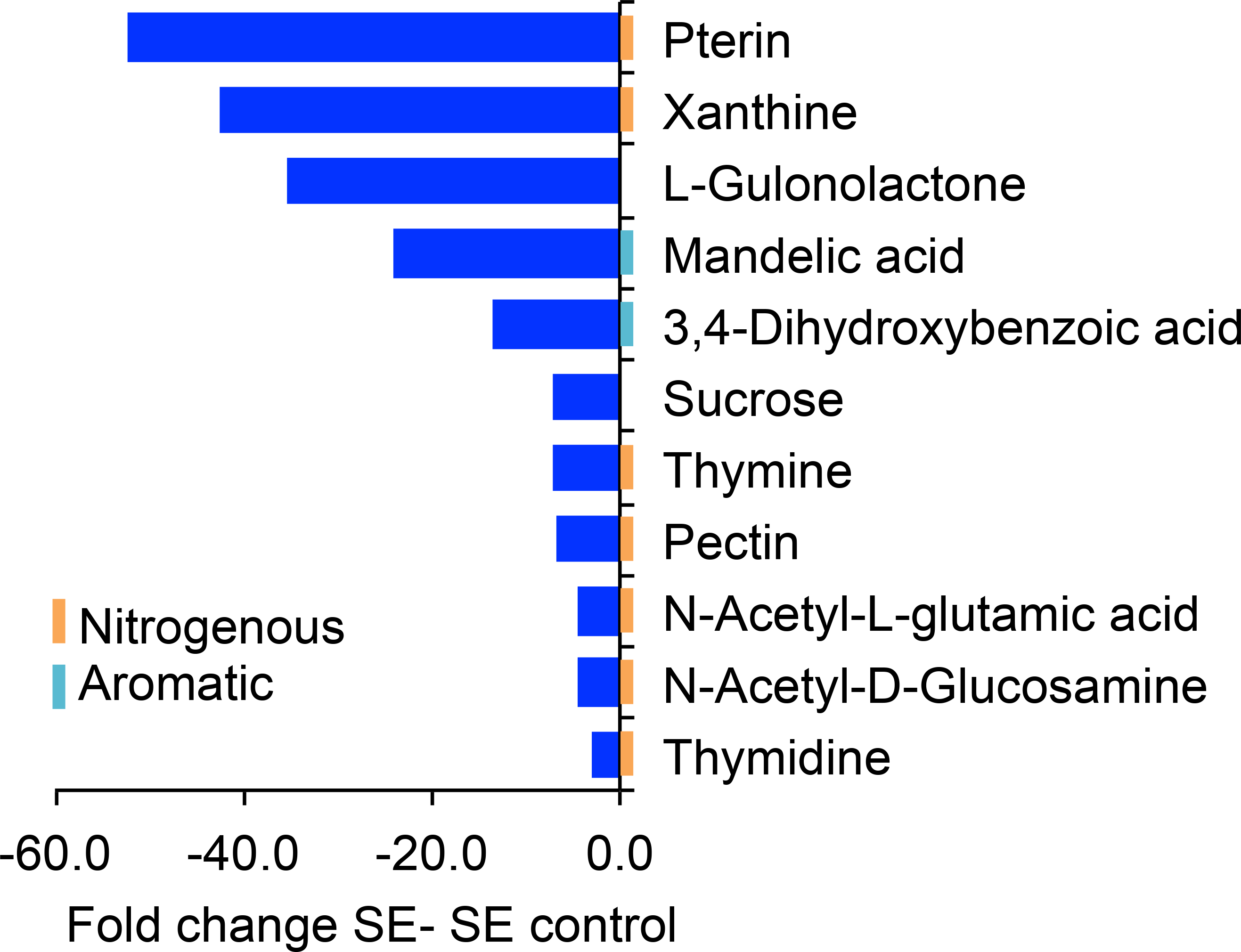
Metabolites reduced in exudates of soil extract grown plants. Fold change of selected metabolites differing between exudates of plants grown in soil extract, and soil extract controls (Anova, p < 0.05). Graphs for single laboratories are given in Fig. S7.

Metabolites that were detected in root tissue and root exudates showed distinct patterns: 42% of these metabolites were significantly different in roots and 43% in exudates, depending on environments. Only 23% of the compounds were significantly different in both datasets, which indicates that root exudates are metabolically distinct from root tissue (Fig. 4, Fig. S5, Table S2, Table S3). We similarly found that only 50% of the metabolites depleted from soil extract were significantly different in root tissues, with 29% of high abundance in soil extract roots (mostly organic acids), 25% of low abundance, and 46% are not detected (nitrogenous compounds).

## Discussion

### Reproducibility of morphological and metabolic data in EcoFABs

This study investigated the reproducibility of morphological and metabolic responses of the model grass *B. distachyon* grown in EcoFABs in phosphate-sufficient and phosphate-deficient mineral medium, and in chemically complex but sterile soil extract. We purposely chose phosphate starvation as an experimental system, as the morphological and metabolic responses of plants are well described, and should be reproducible in a system such as the EcoFAB. The soil extract medium was added to represent a more natural environment, but was sterilized to exclude effects of microbial metabolism on exudation, and to lower variability of the system.

We found that *B. distachyon* fresh weight, phosphate content, and metabolic profiles were distinct for our experimental conditions and that these responses were reproducible across the four participating laboratories. The investigated traits included tissue fresh weight and phosphate content, total root length, and metabolic profiles of roots and exudates. These results compare favorably to a related study comparing three *Arabidopsis thaliana* genotypes grown in soil in pots by ten laboratories (Massonnet *et al.*, 2010) where, similar to this study, materials were distributed from one laboratory, growth conditions were monitored at each laboratory, and one laboratory analyzed leaf morphology, metabolomic and transcriptomic profiles. Although one trait was similar between a core group of four laboratories, all traits significantly varied across laboratories. The authors attributed the variance to the strong influence of small environmental changes in their soil pot system (Massonnet *et al.*, 2010). Our EcoFAB setup comprised a more uniform and controlled growth environment than pots filled with soil, which is likely one cause of the higher reproducibility observed here. Another equalizing factor might have been the use of sterilized soil extract in this study, which did not take into consideration the complex physical and mineral properties of soil, or the effects of microorganisms. It could be that integrating these factors in future EcoFAB studies might increase the variability of the system. It will be important to investigate the reproducibility as well as the morphological and metabolic responses of plants to microbial communities and soil mineralogy, as natural soils were identified as main contributors shaping root morphology, plant carbon exudation, plant-microbe interactions, and rhizosphere extension (Bulgarelli *et al.*, 2013; Koebernick *et al.*, 2017; Holz *et al.*, 2017; Edwards *et al.*, 2018). Overall, we conclude that the reproducibility of plant traits in soil extract EcoFABs is a promising first step towards developing plant growth systems generating reproducible data that are relevant to field environments.

### Metabolic profiles of roots were more distinct than of exudates

Root metabolic profiles were clearly distinct between experimental treatments. Phosphate-sufficient roots were abundant in nucleosides, amino acids, organic acids, and phosphorous compounds, whereas phosphate-deficient roots accumulated nitrogenous compounds, and soil extract-grown roots were deficient in nitrogenous compounds, but accumulated carbohydrates (Fig 4). It will be interesting to investigate if shoot metabolic profiles are similarly distinct between experimental treatments in a future study.

The metabolites detected in *B. distachyon* root exudates in this study (Table S3) were comparable to metabolites detected in exudates of other grasses such as wheat (Iannucci *et al.*, 2017), maize (Carvalhais *et al.*, 2011), rice (Bacilio-Jiménez *et al.*, 2003), *Avena barbata* (Zhalnina *et al.*, 2018) and dicots such as Arabidopsis (Chaparro *et al.*, 2013). Similarly, the *B. distachyon* exudation profile varied with developmental stage, as reported for other plants (Fig. S1) (Chaparro *et al.*, 2013; Zhalnina *et al.*, 2018).

The largest exudate metabolic differences in this study were observed between plants grown in soil extract and soil extract controls without plants. Surprisingly, we did not find many statistical differences in exudates of plants grown in phosphate-sufficient vs. -deficient conditions. For many plants, an increase in organic acid exudation in low phosphate conditions was reported (Neumann & Martinoia, 2002; Plaxton & Tran, 2011; Thijs *et al.*, 2016), which was not found in our dataset. This might be due several reasons. First, plants were grown without phosphate for the entire growth period and might have ceased differential exudation when sampled after three weeks. Second, the small EcoFAB volume likely allows for re-uptake of exuded metabolites, mimicking differential exudation of compounds. Third, the exudation response of *B. distachyon* to phosphate starvation might not be as pronounced as in other species, and be below the detection limit in our assay. Future experiments focusing on the timing and magnitude of *B. distachyon* exudation changes in response to phosphate starvation would be able to address these points. The clear differences observed for fresh weight, tissue phosphate content, and root metabolic profile indicate that the plants indeed were starved for phosphate in our experimental setup.

### Plants deplete metabolites from soil extract

The main differences in exudate metabolic profiles in this study were due to a depletion of metabolites from soil extract by plants (Fig. 5, Table S3). With our experimental setup, we are unable to determine if metabolites are depleted due to uptake by plant roots, or due to for example chemical reactions caused by an altered pH around plant roots. Experiments with isotopically labeled compounds spiked into soil extract could address the fate of metabolites of interest in future experiments.

In addition to depletion of metabolites, a trend for increased citric acid levels in soil extract-grown plants was observed. This might constitute a starvation response, given that exudation of organic acids is a characteristic of phosphate-limited plants (Neumann & Martinoia, 2002; Plaxton & Tran, 2011). The fact that half of the soil extract metabolites, among them organic acids, amino acids, nucleosides, and carbohydrates, are depleted by plants is surprising, as it suggests that plants not only are producers, but also consumers of a significant amount of compounds. Various nitrogenous compounds are depleted from soil extract by plants. Among them is pterin, which is a folate precursor. Folate is an essential part of human diet, and thus, studying uptake of pterin by plants to elevate folate levels might be an interesting biofortification stratey (Strobbe & Van Der Straeten, 2017). Xanthine is part of the purine degradation pathway in plants, and can act as a sole nitrogen source for Arabidopsis thaliana growth (Brychkova *et al.*, 2008). Similarly, thymine thymidine, and N-acetyl-L-glutamic acid could be utilized directly for synthesis of nucleic acids and amino acids, respectively. In addition, plants deplete complex organoheterocyclic compounds such as the ascorbic acid precursor gulonolactone (Smirnoff, 2018), as well as simple carbohydrates such as sucrose. Uptake of these compounds by roots would indicate that plants grow partially heterotrophic in specific environments, importing simple and complex biomass precursors.

There is only a small amount of literature regarding uptake of metabolites by roots: amino acids and sugars were reported to be imported by roots in mineral medium assays where compounds were spiked in (Jones & Darrah, 1994; Yamada *et al.*, 2011), whereas organic acids are likely not imported at significant amounts (Jones & Darrah, 1995). There is evidence that plants are capable to (re)import carbon from environments (Jones & Darrah, 1993), but overall, the scope of how much and which metabolites are taken up by plants from natural environments is currently unknown. In another experimental system comprising of a cyanobacteria and associated heterotrophs, it was found that the primary producer depleted 26% of biological soil crust metabolites, whereas soil heterotrophs only depleted 13% of metabolites (Baran *et al.*, 2015). This might suggest that photoautotroph organisms in general not only release, but also deplete a significant amount of compounds from the environment. Plants might compete with microbes for nutrient soil organic compounds in certain environmental conditions. Besides nutritional functions, compounds could act as signals, as exemplified by a recent study that found the depletion of plant-derived phenolic acids to be associated with rhizosphere microbes (Zhalnina *et al.*, 2018).

Many of the plant-depleted metabolites contained nitrogen, phosphate, or sulfur groups (Fig. 5, Table S3), which suggests that plants not only use inorganic forms, but also more complex compounds as nutrients. Consistent with this hypothesis, compounds containing the N, P, and S groups are low in soil extract-grown roots, likely indicating a fast turnover rate. It was suggested that amino acid uptake might account for 30%-90% of imported nitrogen, depending on the environmental conditions (Jones & Darrah, 1994; Yamada *et al.*, 2011), but overall, data on how much elements are taken up as inorganic vs. organic compounds is missing. In contrast to N, P, and S containing compounds, carbohydrate-type compounds were of high abundance in soil extract-grown roots, likely due to a low external demand for carbohydrates by plant tissues (Fig. 4).

Interestingly, plants depleted metabolites from soil extract in a selective manner, suggesting that the plant controls depletion of metabolites to a certain degree. Similarly, the difference between root and exudate metabolic profiles (Fig. 4, Fig. S5) indicates that plants control exudation to some degree. Selectivity in import and export processes could be achieved by the presence of transport proteins that were described for a number of metabolites (Sasse *et al.*, 2018), and investigation of transport processes is a promising direction for future studies. We conclude that plants not only significantly alter their environment by export, but also by depletion of metabolites.

### Distinct plant growth in soil extract

In this study, plants were grown in basal salt medium widely used in standard laboratory settings, and in soil extract medium that includes water soluble metabolites, but that excludes additional factors defining soils, such as presence of other metabolically active organisms, or solid soil particles.

We observed increased root:shoot ratio in plants grown in soil extract, which might point to nutrient limitations (Cai *et al.*, 2011), consistent with the low phosphate content of soil extract, and of soil extract-grown plants (Fig. 2b, Fig. S2). Interestingly, altered root:shoot ratios were recently also detected for wheat genotypes grown in different soils (Iannucci *et al.*, 2017), suggesting that different soils might affect root:shoot ratio and possibly also metabolic profiles in different ways.

The most prominent phenotypic difference observed for soil extract-grown plants was the four-fold increase in root hair length compared to other plants (Fig. 3). Root hair elongation can be caused by altered nutrient levels (e.g. phosphate, nitrogen, potassium, iron, micronutrients) (Senga *et al.*, 2017; Zhang *et al.*, 2018), and depends on the growth condition used (Nestler *et al.*, 2016). Further, the response to phosphate is concentration dependent (Bates & Lynch, 1996), which might be the cause for the different root hair phenotype observed in phosphate-deficient medium versus phosphate-limited soil extract. Alternatively, the presence of microbes and microbe-derived metabolites that alter plant hormone homeostasis could also cause the phenotype observed in soil extract (López-Bucio *et al.*, 2007; Ortiz-Castro *et al.*, 2011; Zamioudis *et al.*, 2013; Vacheron *et al.*, 2013). Compounds such as tryptophan and salicylate detected in soil extract are reported to alter root morphology (Vacheron *et al.*, 2013), and thus are candidates for causing elongated root hairs. We suggest that the long root hair phenotype observed could be a result of soil extract nutrient levels and specific concentrations of signaling compounds. The determination of the causal factor(s) resulting in the long root hair phenotype represents an important future direction.

Root hair length was shown to have a significant impact on how plants grow in natural soils, and how plants interact with their environment. Root hairs alter physical properties of the soil, such as the extension of the rhizosphere, and the pore size development in soils (Koebernick *et al.*, 2017; Holz *et al.*, 2017). Root hairs also affect biotic interactions by defining the rhizosphere and the amount of carbon exuded from roots (Koebernick *et al.*, 2017; Holz *et al.*, 2017). The complex morphological and metabolic alterations of *B. distachyon* when grown in soil extract stresses the importance of not only considering standard laboratory growth media, but also more natural substrates when studying plant - environment interactions. It would be interesting to investigate how root hair length changes when solid particles, microbial communities, or both are added back to the soil extract used in this study, to investigate morphology changes in a more natural environment. In addition, the observation that increased root hair length was restricted to primary roots but not observed on lateral roots highlights the need for high spatial resolution when measuring root traits, even in a simplified system like the EcoFAB.

In conclusion, EcoFABs are reproducible tools to study a variety of topics, and this reproducibility enables inter-laboratory studies of plant - environment interactions. Their low cost, flexibility, and compatibility with metabolomics studies enables investigations of increasingly complex conditions simulating specific natural environments. We found that *B. distachyon* growth in EcoFABs was reproducible across four laboratories for a number of morphological and metabolic traits, including tissue fresh weight and phosphate content, total root length, and metabolic profiles of root tissue and root exudates. In addition, plants grown in soil extract exhibited an altered root:shoot ratio and elongated root hairs, and depleted half of the investigated metabolites from soil extract. An important next step in the development of more field relevant EcoFABs will be the ability to include solid materials and microbial communities that reflect additional important aspects of soils.

## Acknowledgements

This work was supported by the Microbial Community Analysis and Functional Evaluation in Soils Program at Lawrence Berkeley National Laboratory supported by the U.S. Department of Energy, Office of Science, Office of Biological & Environmental Research, Genomic Sciences Program under contract DE-AC02-05CH11231 at the U.S. Department of Energy Joint Genome Institute, a DOE Office of Science User Facility. The research used resources of the National Energy Research Scientific Computing Center, a DOE Office of Science User Facility supported by the Office of Science of the U.S. Department of Energy under contract number DE-AC02-05CH11231. In addition, J.S. was supported by a NSF grant to University of California Berkeley (NSF Proposal 1617020), and K.Z. by the DE-SC0014079 award to the University of California Berkeley. J.L.D. was supported by NSF INSPIRE grant IOS-1343020, by the Office of Science (BER), US Department of Energy, grant no. DE-SC0014395 and by the HHMI. J.L.D. is an Investigator of HHMI.

## Author contributions

J.S, J.G., and T.R.N. developed the hypothesis, J.S., J.K., B.J.C., A.P.K., J.G., K.L., K.Z. and B.A. conducted experiments, J.S., J.K., B.J.C., A.P.K., B.A., P.S., S.K., B.P.B, K.Z. and D.T. performed data analyses. J.S. and T.N. wrote the paper, and all authors provided comments on the manuscript.

## Supporting information

**Fig. S1** Root morphology and exudate analysis capabilities of EcoFABs

**Fig. S2** Root:shoot ratio of EcoFAB-grown *B. distachyon*

**Fig. S3** Hierarchical clustering of root tissue and exudate metabolites

**Fig. S4** Total root length by laboratory

**Fig. S5** Characteristic metabolites detected in exudates

**Fig. S6** Principal component analysis of soil extract exudate metabolites versus control

**Fig. S7** Metabolites reduced in exudates of soil extract grown plants by laboratory

**Table S1** Participating laboratories and documented growth conditions for the reproducibility experiment

### Excel files

**Table S2** Root tissue metabolite data

**Table S3** Root exudate metabolite data

